# Ecological Relationships in the Serengeti National Park

**DOI:** 10.1101/2020.01.06.895409

**Authors:** Kole Brownstein

## Abstract

The use of camera traps in animal ecology has transformed the field by allowing a greater quantity of detailed observations with limited human interference. One of largest camera trap studies published to date is from the Serengeti National Park (SNP) in Tanzania, East Africa which deployed 225 camera traps and obtained over 1.2 million pictures. This paper will focus primarily on the top predator of the Serengeti, the lion, and how it affects and is affected by prey, subordinate predators and the dramatic shift in the seasons. I asked the following three questions to better understand species relationships in the SNP: 1) Do the seasons in the SNP have an effect on the RAB of lions? 2) Does the presence or absence of lions have an effect on the RAB of hyenas in the wet season? 3) Does the presence/absence of Thompson gazelles (*Eudorcas thomsonii*) and impalas (*Aepyceros melampus*) in the wet season have an effect on the RAB of lions? During the dry and wet season of 2012 in the SNP there did not appear to be a change in the relative abundance of lions (Fig. 1), nor did lion relative abundance affect hyenas. The lions might possibly alter their diet during the dry seasons to include non-migratory species and choose not to change their territory on a season to season basis. Based on these findings research should focus on how the lions adapt to the changes in prey abundance during the wet and dry season. This preliminary analysis of the ecological dynamics of lions and associated species in the SNP is intriguing and yet raises significantly more questions than it has answered. Additional research surrounding the effects of yearly migration on the lions home territory, diet, and species interactions should be investigated more thoroughly, to greater understand the ecological relationships in the SNP.

## Introduction

The use of camera traps in animal ecology has transformed the field by allowing a greater quantity of detailed observations with limited human interference, leading to significant discoveries in ecological relationships and population dynamics between species (O’Connell et al. 2010). Camera traps allow for a nonintrusive approach to viewing and studying animal behavior for even the most cryptic species, such as Himalayan snow leopards (*Panthera uncia*) (O’Connell et al. 2010) or honey badgers (*Mellivora capensis*) (Allen et al. 2018) which are nearly impossible to see or document. One of largest camera trap studies published to date is from the Serengeti National Park (SNP) in Tanzania, East Africa (Swanson et al. 2015), which deployed 225 camera traps over 1,125km^2^ and obtained over 1.2 million pictures across 99,241 camera trap days (Swanson et al. 2015), and are continuing to collect data. Due to the extensive digital collection of photographs, the researchers used the general public to classify each image, which was then compiled into a massive dataset after an algorithm was used to ensure appropriate and correct classifications (Swanson et al. 2015). The specific focus of this study was to examine and “evaluate spatial and temporal dynamics of large predators and their prey” (Swanson et al. 2015). Specifically, the SNP has been prominent study area since the 1960s by the Serengeti Lion Project to assess population and range sizes of lions (*Panthera leo*) and this data set has allowed for a multitude of additional research to be conducted in the region (Swanson et al. 2015).

Predator-prey relationships are highly complex and camera trapping has allowed for an extensive and in-depth look at how species respond to one another. Instances of predation, interference, and movement are connected to and driven by a need for “resource acquisition” (Sinclair and Arcese 1995). Each organism has fundamental requirements such as food, shelter, mating, and most importantly to avoid becoming a meal (Sinclair and Arcese 1995). While maintaining these fundamental needs organisms will disrupt and alter others for their individual success, creating a variety of downstream trophic effects (Sinclair and Arcese 1995). From a wide-angle view, changes in rainfall in the dry and wet seasons in the Serengeti change the availability of grazing lands which causes massive migrations including 1.6 million zebras and wildebeests (Swanson et al. 2015). Due to the changes in prey abundance between seasons, camera traps can be used to evaluate the predatory abundance and more specifically look at how prey and predatory abundance is affected by one another. The extensive nature of the photobank and analysis has opened the door to truly examine the population and behavioral dynamics of the species who inhabit the SNP. However, this paper will focus primarily on the top predator of the Serengeti, the lion, and how it affects and is affected by prey, subordinate predators and the dramatic shift in the seasons.

In the Serengeti, the lions are the top predators followed by subordinate predators such as the hyena (*Crocuta crocuta*) and cheetahs (*Acinonyx jubatus*). Top predators will usually “kill, harass, and steal food from less dominant predators” creating a clear and evident hierarchy (Swanson et al. 2016). These subordinate predators will avoid high-intensity interactions and find other means to scavenge and hunt that do not elicit a response from in this case the lions (Swanson et al. 2016). The term “landscape of fear” coined by Laundré et al. (2001), describes instances “in which animals, in their effort to reduce their vulnerability to predation… have increased their levels of vigilance” (Laundre et al. 2001). This term was used to describe how gray wolves (*Canis lupus*) in Yellowstone National Park created an environment of higher vigilance in prey and now it can be used to discuss the relationship between subordinate predators and prey of the lions in the Serengeti (Swanson et al. 2016, Laundre et al. 2001). Subordinate predators will typically stay away from lion prime areas of activity and potentially avoid locations where there are higher levels of food, water, and shelter (Swanson et al. 2016). More specifically to the interactions in the Serengeti, research shows that hyenas and lions share similar diets and will intimidate one another as well as steal food (Swanson et al. 2016). In most scenarios, the lions maintain dominance over the hyenas through aggressive attacks and stealing, but when hyena pack sizes are large enough, they can overpower lions (Swanson et al. 2016). However, these occasions of subordinate aggressive maneuvers to steal food from the lions do “not appear to inflict a measurable cost to the lions” (Swanson et al. 2016).

Based on the findings of Swanson et al. 2016 with the “landscape of fear” there is also an effect on the lion’s prey as well. As mentioned above the presence of the wolves in Yellowstone caused increased vigilance in wolf prey (elk, *Cervus elaphus*, and bison, *Bison bison*) and thus it would be probable that prey species of the lions would also have heightened vigilance and avoid areas where lions are present (Swanson et al. 2016). In addition, there is the potential to see decreases in feeding due to increases in vigilance, and prey will be more particular about the general landscapes they feed on in an effort to avoid predation (Swanson et al. 2016). The relationship between predator and prey can also be examined in the reverse. Due to the changes in prey movement to avoid predation top predators could potentially move into areas with a higher abundance of prey. Taking the Yellowstone example, wolves might venture out of their regular territory to target prey that is not at a heightened vigilant state. There are clearly many potential interactors that can alter and mitigate the behavior of lions, hyenas, and their prey.

As this data set is immense and robust, I have chosen to focus on a small set of the information obtained from only January, February, March (wet season) and July, August, September (dry season) during the year 2012. By examining the relative abundance (RAB) of lions and other species throughout the study area I hope to learn how species interact with one another and are affected by the environment. I asked the following three questions to better understand species relationships in the SNP: 1) Do the seasons in the SNP have an effect on the RAB of lions? 2) Does the presence or absence of lions have an effect on the RAB of hyenas in the wet season? 3) Does the presence/absence of Thompson gazelles (*Eudorcas thomsonii*) and impalas (*Aepyceros melampus*) in the wet season have an effect on the RAB of lions?

## Methods

### Study Sight and Image Aggregation

From June of 2010 to May of 2013, Swanson et al. (2015) set up and maintained 225 camera traps in a 1,125km^2^ grid within the 25,000km^2^ Serengeti National Park (Swanson et al. 2015). The camera grid was located inside the “long-term lion study area” which contains a diverse landscape (Swanson et al. 2015). They placed cameras at the center of 5 km^2^ grid cells on the nearest tree that would avoid potential misfires from any movement not associated with an animal (Swanson et al. 2015). In addition, they placed the cameras at height of 50cm with a goal to capture medium-large mammals (Swanson et al. 2015).

During the study period, 1.2 million images were captured by Swanson et al, over the course of close to 99,000 camera trap days (Swanson et al. 2015). In the end, it was determined that slightly over 300,000 of the images were of intended species and the rest were misfires (Swanson et al. 2015). All images were screened multiple times by volunteers to ensure success with classification and standard requirements (Swanson et al. 2015). Following, Swanson et al. (2015) subjected the images to a verifying algorithm that compared the results of the public classifications to determine any instances of error (Swanson et al. 2015). The full data set was then published so that scientists around the world could continue to research species dynamics in the SNP.

### Data Manipulation

I only examined and investigated camera trap events of lions, hyenas, Thompson gazelles and impalas. The original data set is arranged as rows of unique images with columns of additional information for each image including the presence of a species, which I then set up to examine the relative abundance of predators and prey in the SNP. I defined an “event”, based on Allen et al. (2018), as any time a species was seen and removed all triggers/images following the initial event for a 30min period (Allen et al. 2019). Removing all images for a 30 minute period flowing the initial event, reduced the possibility of having the same animal count multiple times in a short time frame (Allen et al. 2019). Following Allen et al. (2018), I then determined the number of events per camera and divided by the number of events per total trap-nights for each camera and calculated the number of events per 100 trap-nights of each camera using the formula:

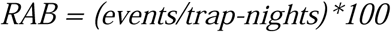

to standardize RAB values as units of events per 100 trap nights.

Following are the methods for the data manipulations and statistical test completed for each of the three questions. Note that all statistical tests, assumption tests, and calculations were completed using the software platform Rstudio.

### Question 1: Lion RAB in Wet and Dry Season

To compare whether the relative abundance of lions in the dry and wet season is different, I attempted to use a 2-Sample t-test. I re-arranged the data from its original set up so that one column contained the lions RAB value which was a numerical continuous variable and the second column contained a categorical variable of seasons with two levels either the “dry” or “wet” season. Following the guidelines of running statistical analyses, I checked the assumptions for the 2-Sample t-test starting with ensuring random sampling which was confirmed by the original research conducted by Swanson et al. 2015. After, I checked to make sure the RAB values for the two levels were normally distributed using the *hist* function in Rstudio. The initial histograms were extensively right-skewed. In an effort to normalize the data I performed natural log, log, and square root transformations. As there were numerous zero points throughout the data set, I completed the above transformation while adding one to each lion RAB value for the log and natural log transformations. Again, the histograms showed right-skewed data indicating that the data would not normalize through transformations. As one of the major assumptions for a 2-Sample t-test was met, a non-parametric test was required. I then followed the procedure to run Wilcoxon Rank Sum test which compares the mean RAB values between the two seasons. The test was a non-parametric test, which required only the assumptions that the data is randomly sampled which has been previously confirmed. I ran the Wilcoxon Rank Sum test and a p-value was obtained.

### Question 2: Hyena RAB in the presence and absence of lions

To investigate whether the presence/absence of lions in the wet season effects on Hyena RAB, I again attempted to use a 2-Sample t-test. I rearranged the data so that I had a column of hyena RAB which was continuous and numerical and a second column containing a categorical variable with two levels, presence (RAB > 0)/absence (RAB = 0) of lions. Similarly, to the first question while the data was randomly sampled the histograms showed that the hyena RAB data was not normally distributed in the presence or absence of lions. As above, I ran the exact same transformations which revealed no changes to the normality of the data. I then choose to run a permutation test, a non-parametric test that can be used compare the means of two groups. The assumptions were met for this test as the data was randomly sampled and the distributions of the two levels (presence and absence of lions) had similar means of 2.99 and 3.09 (Table 2) and both histograms were right skewed and almost identical in shape but not in scale. Following, I ran the permutation test with 10,000 permutations and calculated the p-value.

**Table 1:**
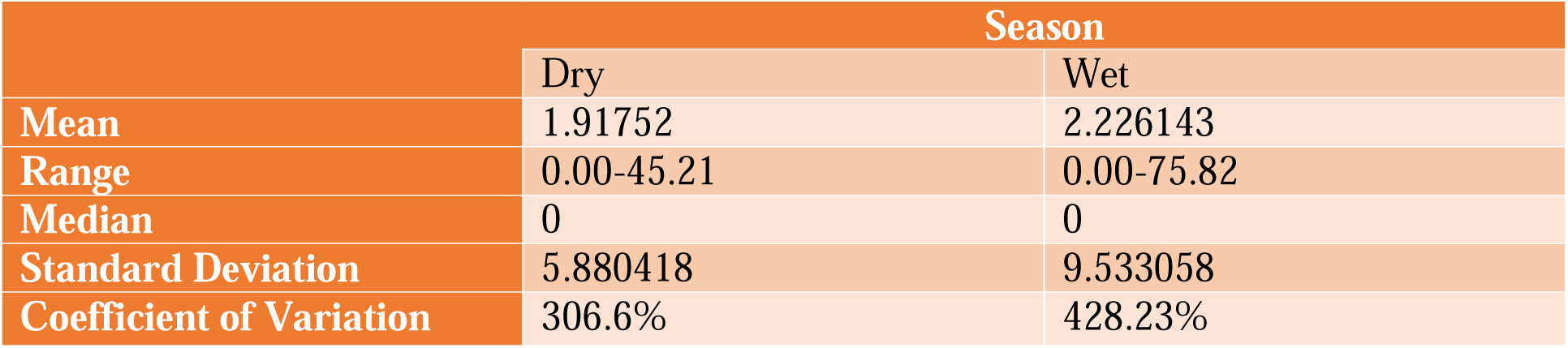
Summary Statistics of Lion RAB in the Dry and Wet Season of 2012.

|  | Season |  |
| --- | --- | --- |
|  | Dry | Wet |
| **Mean** | 1.91752 | 2.226143 |
| **Range** | 0.00-45.21 | 0.00-75.82 |
| **Median** | 0 | 0 |
| **Standard Deviation** | 5.880418 | 9.533058 |
| **Coefficient of Variation** | 306.6% | 428.23% |

**Table 2:** Summary Statistics of Hyena RAB in the presence and absence of lions (wet season 2012)

|  | Absent | Present |
| --- | --- | --- |
| Mean | 3.091881 | 2.986154 |
| Range | 0.00-27.27 | 0.00-29.21 |
| Median | 0 | 1.1 |
| Standard Deviation | 5.300861 | 5.58516 |
| Coefficient of Variation | 171.44% | 187.04% |

### Question 3: Lion RAB in the presence and absence of Thompson gazelles and impalas

To explore the effects of the presence and absence of Thompson gazelles and impalas on the RAB of lions I attempted to run an ANOVA test. I simplified the data so that I had one column of Lion RAB (numerical and continuous) and a second column containing a categorical variable with four levels: absent gazelles and impalas (NGNI), present gazelles and impalas (PGPI), absent gazelles and present impalas (NGPI), present gazelles and absent impalas (PGNI). None of the four levels had a normal distribution in terms of Lion RAB with any of the transformation attempted in question 1 and 2. All histograms were extensively right-skewed. Because the data did not meet the requirements for the ANOVA test, I performed a Kruskal Wallace test, a non-parametric alternative to the ANOVA and determined a p-value.

## Results

During the dry and wet season of 2012 in the SNP there did not appear to be a change in the relative abundance of lions (Fig. 1). In the dry season, RAB ranged from 0.0 to 45.21 with an average of 1.91 whereas in the wet season the RAB ranged from 0.0 to 75.82 with an average of 2.23 (Table 1). Variation in the RAB of lions in the dry season had a standard deviation (SD) of 5.88 events with 306.6% coefficient of variation (CV) and a SD of 9.53 with a CV of 428.23% in the wet season (Table 1). There was no significant difference in lion relative abundance during the dry and wet season of 2012 (W = 9091, p-value = 0.5009).

**Figure 1.**
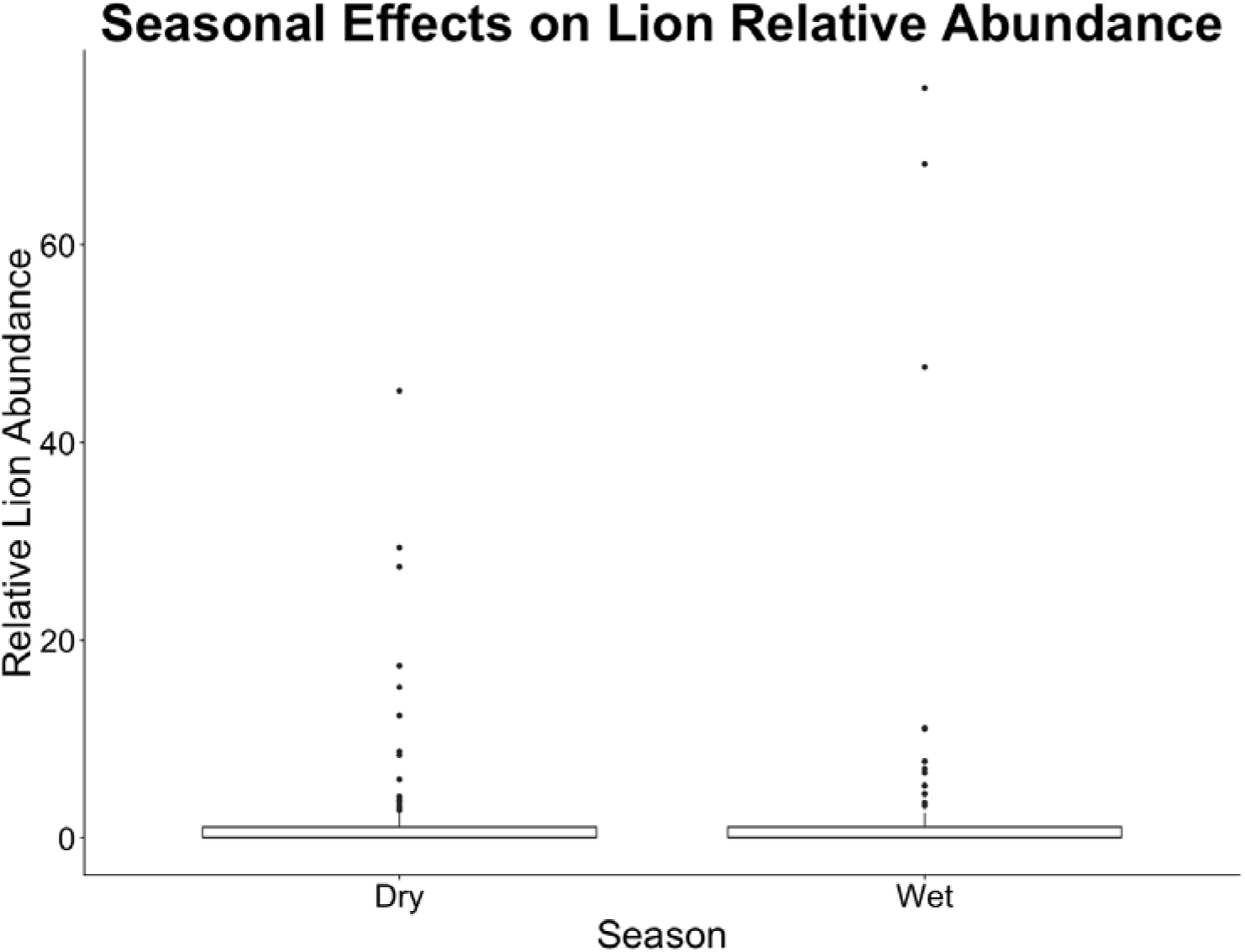
The relative abundance of lions in the dry and wet season in the Serengeti National Park observed by162 camera traps during 2012. Lion relative abundance (RAB) is given in units of the number of events in 100 trap nights. RAB values were determined by using motion censored cameras to identify the presence of a Lion and then disregarding all sightings for a 30-minute period following initial observation. This sighting/identification was referred to as an event. The total number of events for each camera was calculated and divided by the number of trap nights the camera was operational. The value was then multiplied by 100 to give the number of events per 100 trap nights. P-value > 0.05.

During the wet season of 2012, the presence and absence of lions did not have an effect on the RAB of hyenas (Fig. 2). In the absence of lions, hyena RAB ranged from 0.0 to 27.27 with an average of 3.09 whereas when lions were present the RAB ranged from 0.0 to 29.21 with an average of 2.99 (Table 2). Variation in hyena RAB in the absence of lions was found to have a SD of 5.300861 and a CV of 171.44% and in the presence of lions a SD of 5.58 and a CV of 187.04% (Table 2). There was no significant difference in hyena RAB in the wet season of 2012 in the presence or absence of lions (p-value = 0.9626).

**Figure 2.**
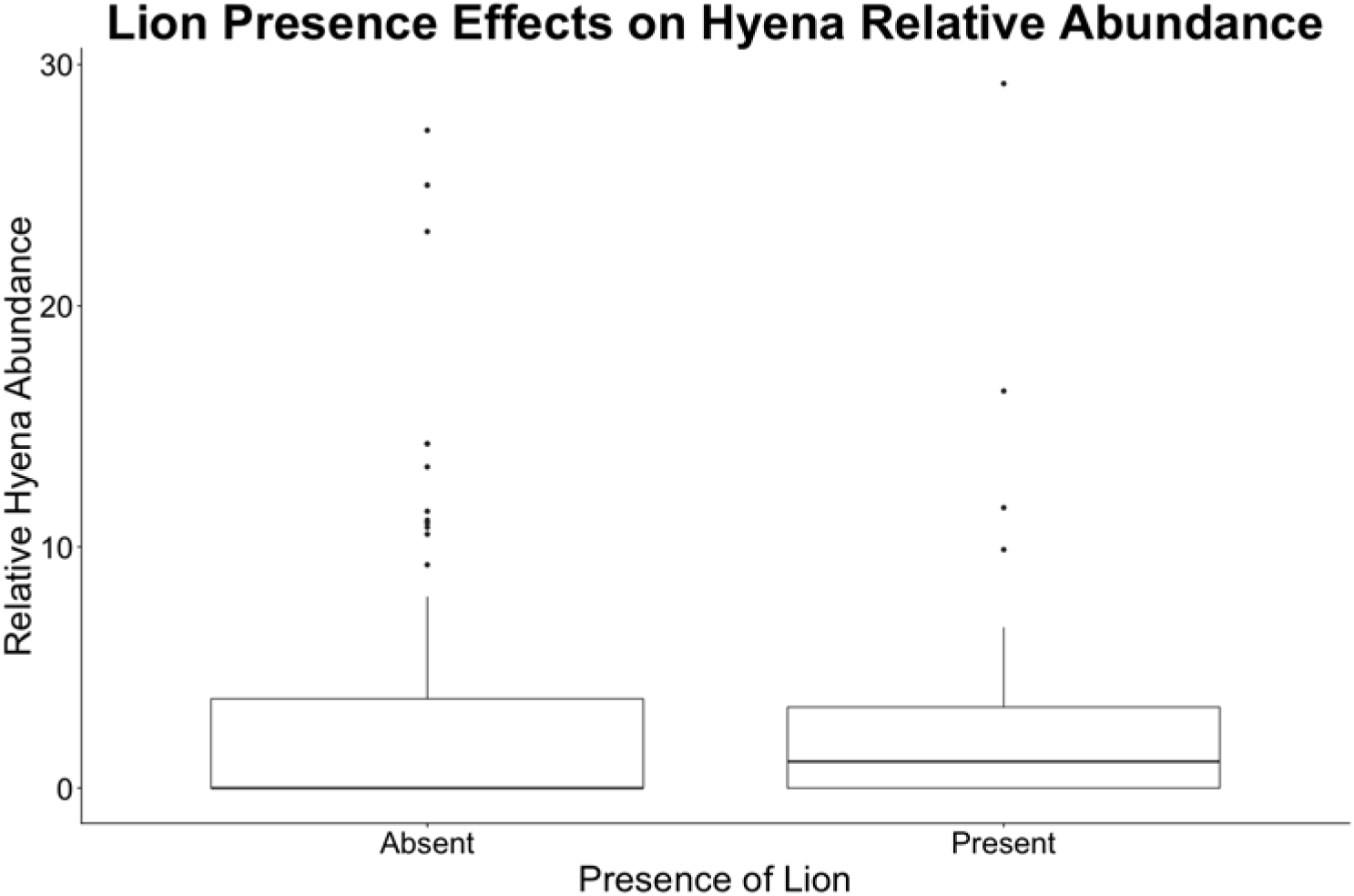
The relative abundance of hyenas in the 2012 wet season (at 140 camera location where lions were either present or absent. Hyena relative abundance (RAB) is in units of events per 100 trap nights. RAB values were determined by using motion censored cameras to identify the presence of a Hyena and then disregarding all sightings for a 30-minute period following the initial observation. This sighting/identification was referred to as an event. The total number of events for each camera was calculated and divided by the number of trap nights the camera was operational. The value was then multiplied by 100 to give the number of events per 100 trap nights. P-value > 0.05.

**Figure 3.**
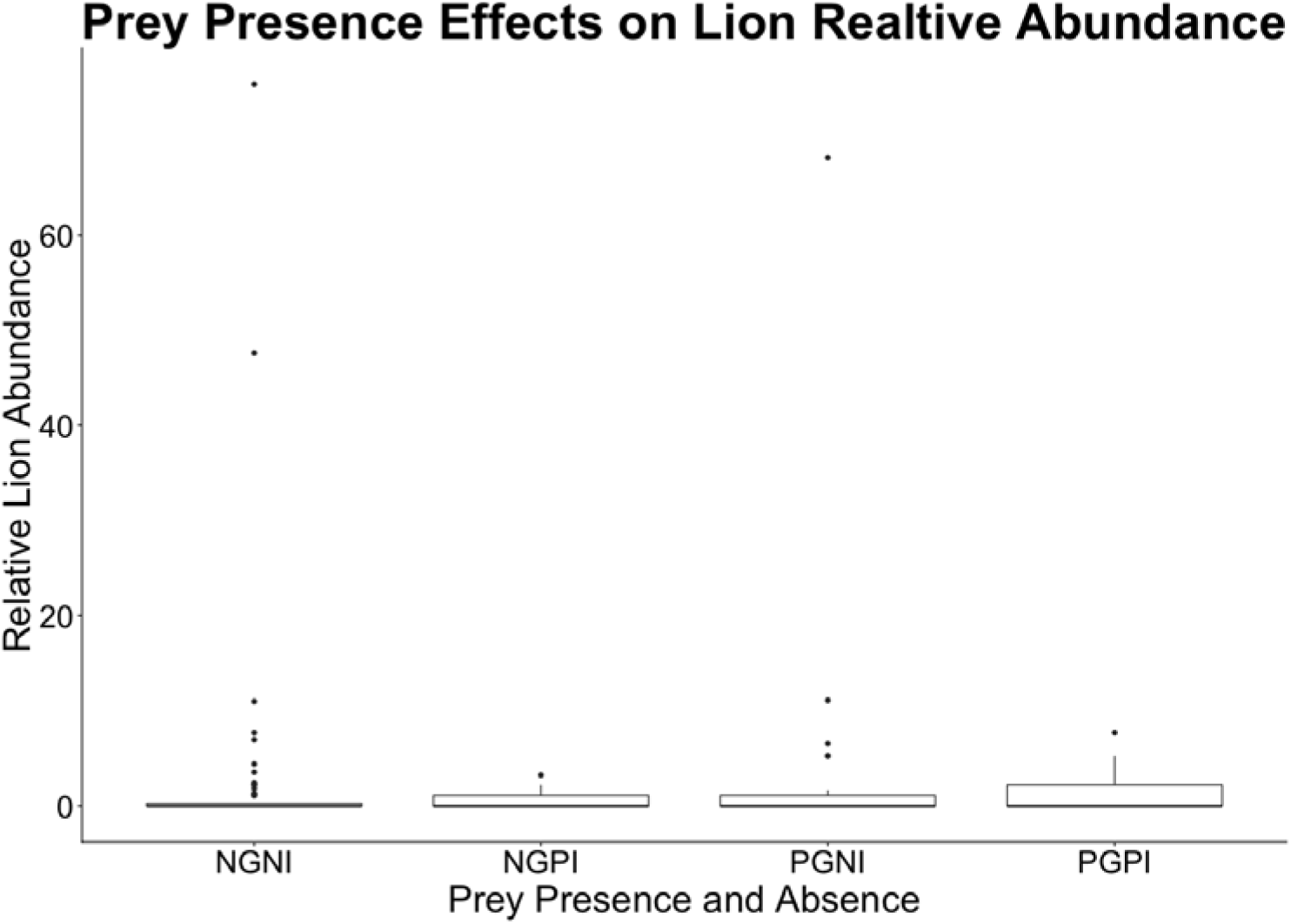
The presence of impalas and Thompson gazelles and the effect on the relative abundance (RAB) of lions during the wet season of 2012 in the SNP. 140 camera sites were used over the three-month period in the wet season. Treatments include: absent gazelles and impalas (NGNI), present gazelles and impalas (PGPI), absent gazelles and present impalas (NGPI), present gazelles and absent impalas (PGNI). Lion RAB is in units of events per 100 trap nights. RAB values were determined by using motion censored cameras to identify the presence of a Lions and then disregarding all sightings for a 30-minute period following the initial observation. This sighting/identification was referred to as an event. The total number of events for each camera was calculated and divided by the number of trap nights the camera was operational. The value was than multiplied by 100 to give the number of events per 100 trap nights. P-value > 0.05

Differing treatments of present and absent prey combinations of Thompson gazelles and impalas had no effect on the level of lion RAB in the wet season of 2012. When both the gazelles and impalas were not present lion RAB had a calculated mean of 2.5, a range of 0 – 75.82 and SD of 10.83 with a CV of 423.64% (Table 3). When gazelles were not present, but impalas were present, lion RAB had a calculated mean of 0.51, a range of 0 – 3.23 and SD of 0.93 with a CV of 180.86% (Table 3). In addition, when both gazelles and impalas were present, lion RAB had a calculated mean of 1.17, a range of 0 – 7.69 and SD of 2.18 with a CV of 186.43% (Table 3). Lastly, when gazelles were present, but impalas were absent, lion RAB had a calculated mean of 3.413, a range of 0 – 68.13 and SD of 12.49 with a CV of 365.86% (Table 3). In the varying combinations of prey presence and absence of both Thompson gazelles and impalas, there was no significant difference in the lion RAB (chi-squared = 0.73867, df = 3, p-value = 0.8641).

**Table 3:** Summary statistics for the present and absent combinations of Thompson gazelles and impalas and their effect on the relative abundance of lions in the wet season of 2012.

|  | NGNI* | NGPI* | PGPI* | PGNI* |
| --- | --- | --- | --- | --- |
| **Mean** | 2.557206 | 0.5142857 | 1.170476 | 3.413 |
| **Range** | 0.00-75.82 | 0.00-3.23 | 0.00-7.69 | 0.00-68.13 |
| **Median** | 0 | 0 | 0 | 0 |
| **Standard Deviation** | 10.83344 | 0.9301482 | 2.182167 | 12.48678 |
| **Coefficient of Variation** | 423.64% | 180.86% | 186.43% | 365.86% |

\*Note: Treatments include: absent gazelles and impalas (NGNI), present gazelles and impalas (PGPI), absent gazelles and present impalas (NGPI), present gazelles and absent impalas (PGNI).

## Discussion

There are many interactions and complex processes that surround the relationships of different animal species. The Serengeti National Park is home to numerous animal species with varying hierarchical dynamics (Sinclair and Arcese 1995). The top predator in the park is the lion with subordinate predators such as the cheetah and hyena and below them are multitudes of sizeable herbivorous prey. As discussed in Swanson et al. (2015) and Laundré et al. (2001) a “landscape of fear” dynamic can be and is formed between species of varying positions in the food web. Prey in close proximity to predators can have elevated instances of vigilance, eat less and choose specific areas to feed with greater visibility than those species, not in predator territories (Swanson et al. 2016). Not only will predator and prey be potentially affected by the landscape of fear but there can also be predator – predator dynamics. As the lion is the top predator it is possible that hyenas will avoid areas of potential conflict and competition and have different territories.

The SNP as discussed above is dramatically affected by the changing of the seasons. Massive migration events take place each year as the zebra, wildebeests, and other species move to richer grazing lands. In the dry seasons, the zebra and wildebeest are expected to be in the northern regions of the SNP, outside the camera trap study area. Whereas in the wet season the zebra and wildebeest are expected to be centrally located in the study area (Boone et al. 2006). Based on the movements of the large prey it was expected that the lions might have lower levels of relative abundance in the study area as they would move dynamically with the herds. The statistical analysis indicated that there was no change in the relative abundance of the lions between the wet and dry season of 2012 (Fig 1). The lions might possibly alter their diet during the dry seasons to include non-migratory species and choose not to change their territory on a season to season basis. Based on these findings research should focus on how the lions adapt to the changes in prey abundance during the wet and dry season. Questions should be asked concerning what do the lion’s diet consist of during the wet season compared to the dry season? Further clarification should be made into determining if the lions adjust home ranges on a seasonal basis in response to migration events by increasing the number of study years.

In relation to the interactions between the lions and the hyenas, it was hypothesized that the hyenas would tend to avoid areas that lions frequently inhabited. However, the test revealed that during the wet season of 2012 the hyena RAB was not affected by any level of presence of lions (Fig 2). As hyena and lions diets are highly similar it makes sense that both species would be near one another due to resources. But based on the landscape of fear argument of Swanson et al. (2016), hyenas would avoid areas where lions were present to avoid any impending conflicts leading to injury or death. Hyena pack sizes might be large enough that they are in fact able to cohabitate and compete with the lions and thus not afraid or concerned with any potential conflict and competition. In regard to the effect of lions on hyenas, research should be expanded to examine the species interactions in depth and compare RAB levels in the wet and dry seasons. It is possible that the dynamics between two species change when the availability of prey diminishes due to the migration events and more typical behaviors between species would be observed.

I hypothesized that prey abundance would be indicative of predator abundance. However, during the wet season of 2012 the presence or absence of impalas and Thompson gazelles had no effect on the relative abundance of lions. Generally it was expected that with the presence of both prey species there would be an increase in lion relative abundance but in fact, no combination of either species indicated a significant difference. Further research should be conducted to further investigate if there are additional prey that can affect lion RAB. Only two prey species were examined in this analysis and the Swanson et al. (2015) data set includes numerous other prey species. In addition, comparisons on prey presence and its effects on lion RAB should be made in both the dry and wet season. This could indicate the specific species that influence and predict lion territories in both of the seasons with the varying stressors.

Overall, in future studies I would incorporate additional years of data. Currently, only 2012 is represented in this analysis and expanding the time frame will increase the number of data points making the data set more robust and predictive of the future events. This preliminary analysis of the ecological dynamics of lions and associated species in the SNP is intriguing and yet raises significantly more questions than it has answered. Additional research surrounding the effects of yearly migration on the lions home territory, diet, and species interactions should be investigated more thoroughly, to greater understand the ecological relationships in the SNP.

